# Wealth-relative effects in cooperation games

**DOI:** 10.1101/333971

**Authors:** Robert L. Shuler

## Abstract

This paper investigates conditions under which game agents benefit from considering wealth relative to decision payoff, presents simulation analysis of these effects, and explains why they often do not show up but it is realistic that they should. We extend the known categories of games reported to exhibit wealth relative effects (chicken games) to many others (including Prisoner’s Dilemma) while clarifying that the poor must avoid survival risk, regardless of whether this is associated with cooperation or defection. A simulation of iterated Prisoner’s Dilemma with wealth accumulation and a survival threshold (which we call the Farmer’s Game) is used to evaluate tit-for-tat and four variants, including Subsist, Thief, Exploit and Middle (even lower risk than Subsist). Equilibrium payoffs are used to keep the game scaled to social relevance, with a fraction of all payoffs externalized as a turn cost parameter. Findings include poor performance of tit-for-tat near the survival threshold, superior performance of Subsist and Middle for both poor and wealthy players, dependence of survival of the poor near the threshold on tit-for-tat forgiveness, unexpected optimization of forgiveness without encountering a social dilemma, improved performance of a diverse mix of strategies, and a more abrupt threshold of social catastrophe for the better performing mix. Additionally we find that experimental results which appear to be at odds with conventional findings of cooperation vs. network size can be reconciled with theory and simulation via wealth-relative weighting, which opens the door to practical application of cooperation theory.

**Significance Statement:** Enabling comparison of theoretical and simulated game cooperation theory results to controlled experiments with live subjects and in-situ data from field surveys will enable application of scientifically verified results to societal and policy problems, and will generate new and unexpected insights through clearer interpretation of data. Extension of wealth-relative effects to a broader range of games also allows analysis of real life situations with greater confidence.

## Introduction

Games with a predetermined payoff matrix are frequently used to study the formation, maintenance or evolution of cooperation. The most common example is Prisoner’s Dilemma, though many others are used. The games can be played on a one-shot or iterated basis, pair wise or multi-player. Frequently the dependency of results on network relations among participants is studied. For large network and evolutionary studies a simulation approach may be used rather than actual participants, but live subjects are also recruited and their behavior studied.

It has been found that in the general case cooperation develops under limited conditions such as kin or group selection (Hamilton 1964; Maynard Smith 1964; Wilson 1974; Wilson and Wilson 2007; Bowles and Gintis 2011; Simon et. al. 2012). Reciprocal cooperation among non-kin in excess of purely rational behavior (Nash Equilibrium) can develop in evolutionary settings without kin/group selection (Trivers 1971; Axelrod and Hamilton 1981). Some instinct for cooperation is evident from early emergence in infants (Tomasello and Gonzalez-Cabrera 2017), as well as an instinct for fairness (Burns and Sommerville 2014). Culture, especially its co-evolution with cooperation, plays an important role as well (Boyd and Richerson 2009). Social learning likely plays a role in the formation of patterns on which cooperation or conflict may be based (Pulliam 1982), though recent evidence suggests that direct social learning of cooperation strategies may guide behavior toward individual utility maximization strategies at the expense of costly cooperation (Burton-Chellew, Mouden and West 2017).

## Problem Description

This paper first addresses the general problem of experimental confirmation of the results of research into the evolution and modeling of cooperation using game theory. We proceed by examining and relating data from two papers, one conducting experiments with live participants, the other based on simulations using both real networks and randomized modifications of those networks.

The hypothesis needed to reconcile the experimental and empirical results is based on a narrow analysis applicable to chicken games which presumed an accumulation of wealth or fitness from game playing, but chicken games characterize neither the empirical nor experimental data used. Thus in a subsequent section we explore more broadly how this hypothesis might apply using simulation, and find the conditions under which it does apply to Prisoner’s Dilemma but only under conditions approximating social persistence such as we might find in real life, and which simple integer idealized games quickly remove. The tailored version of Prisoner’s Dilemma used we call the Farmer’s Game. Several important corollary observations emerge from the simulations.

Game theory frames decision problems in an exact but limited way so that concepts are clear and theorems can be proved. It is implicit that the game in question is a complete description of the options available to the players; otherwise they may elect not even to play but to make investments elsewhere. It is also assumed that only the magnitudes of payoffs relative to the other payoffs are relevant. Experiments with human subjects rarely meet these criteria.

Since defining the complete environment of test subjects as a game is usually infeasible, then comparison among simulated, experimental and in-situ data is problematic. Without such comparison, cooperation theory based on game theory can hardly make definite analyses of real world problems, except in special cases where participants either behave as if the subject game were critically important, or their existence is literally dependent on it and little else. That includes other factors in the environment, and any stored payoff (wealth or fitness) they may have. For these reasons game/cooperation theory has been remarkably successful in some areas while inexplicably failing in others, the failures usually attributed ad hoc to non-rational behavior.

There has been some success with evolutionary simulation approaches in predicting not strictly rational behavior. Non-kin reciprocal cooperation has been found to develop and to depend on either small group size (Maynard Smith 1976; Killingback, Bieri and Flatt 2006), or heterogeneous or Small-World networks (Pacheco and Santos 2005) so that clusters of cooperators can fend off defectors (Lozano, Arenas and Sánchez 2008). Defectors, in this context, make purely rational choices. Persistent links in a large network have been found to be sufficient (Axelrod, Riolo and Cohen 2002).

Lozano, Arenas and Sánchez (2008) studied two networks from the real world using simulation and Prisoner’s Dilemma, considering both the actual network and a randomized version of each network which preserved node degree but not clustering. One was an EMAIL network with a high degree of connectivity within and between communities, and the other a PGP (pretty good privacy) network that was more hierarchical. The real EMAIL network and the randomized versions of both networks had similar relations between the temptation parameter *b*, starting with 0.95 density of cooperators in both randomized cases, to 1.0 density of cooperators in the real EMAIL network for *b=1* (no temptation, defection payoff same size as cooperation payoff). Density decreased slowly at first and then more rapidly to a range of 0.15 to 0.3 for *b=2* (defection payoff double the cooperation payoff).

Figure 1 shows the real and randomized simulation results from Lozano et. al. for both networks. The real world PGP network performed differently, holding a lower level of initial cooperation relatively stable over a broad range of temptation before finally collapsing near *b=2*. The randomized PGP network was very similar to the randomized EMAIL network, illustrating Lozano’s main point that global network parameters are not the most important factors in cooperation. A second point we could take from Lozano et. al. may be that the relation between temptation and cooperation is similar over a broad range of networks unless they have very particular mid-scale structure (mezoscopic is Lozano’s term).

**Fig. 1.**
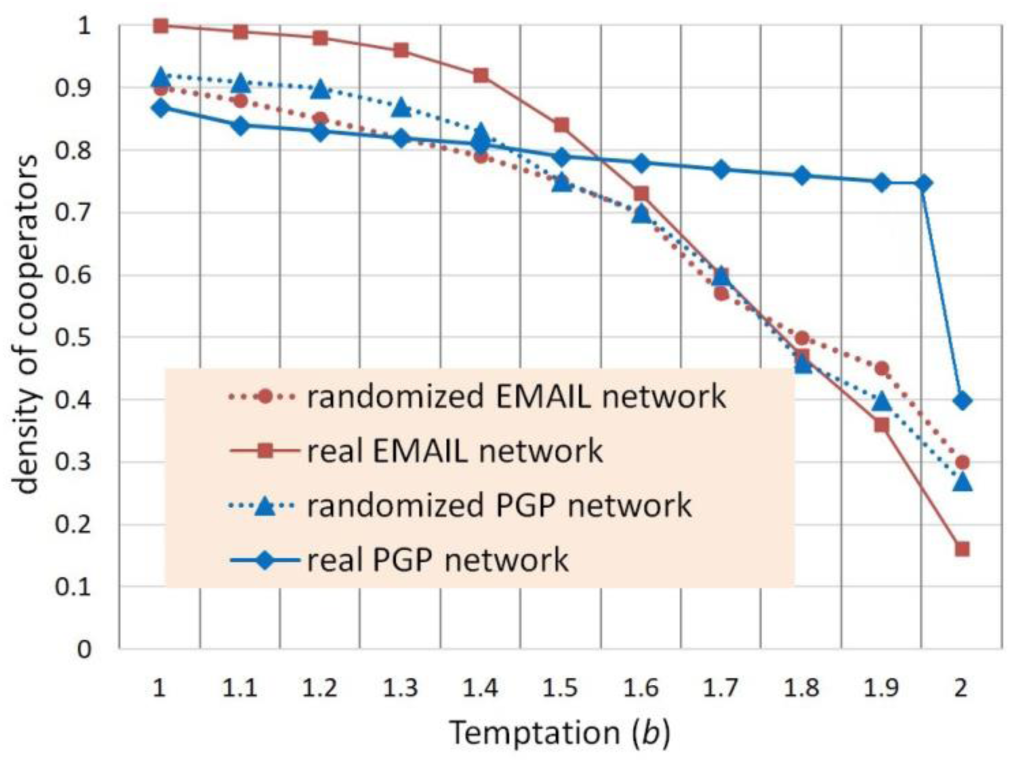
Density of cooperators from Lozano et. al. 2008.

Other investigators such as Xu, Wang, Deng and Li (2014) report roughly similar results for cooperation as a function of temptation, even for simple lattice networks, with cooperation always vanishing below *b=2*.

Buchan et. al. (2009) report on an experiment with live subjects from six different countries designed to evaluate the effects of the networks and cultural attitudes of the participants’ home countries, over a range of degrees of “globalization” which they describe as “*the increased connectivity and interdependence among people worldwide*.” The network of the experiment was a hierarchy of three LOCAL groups with four INDIVIDUALs in each, all comprising a WORLD. There were only three trials, presumably to limit social learning during the experiment, since the objective was to measure effects of the home networks, not the experimental network.

In each trial participants were given 10 tokens worth $0.50 USD each and asked to allocate them to INDIVIDUAL, LOCAL or WORLD accounts. The amount at risk over the three trials is then $15 per participant. For non-USA participants, tokens represented equivalent purchasing power in their home countries. Experimenters doubled the amounts in LOCAL accounts, dividing the proceeds among participants in the local group, and tripled the amount in the WORLD account dividing it among all participants. Thus the collective optimum would be for all participants to contribute all tokens to the WORLD account, while the Nash Equilibrium would be to invest only in the INDIVIDUAL account.

For our purposes we sum the amount contributed to LOCAL or WORLD accounts and normalize as a percentage for an indicator of the extent of cooperation, shown as the green dot-dash line in Figure 2.

**Fig. 2.**
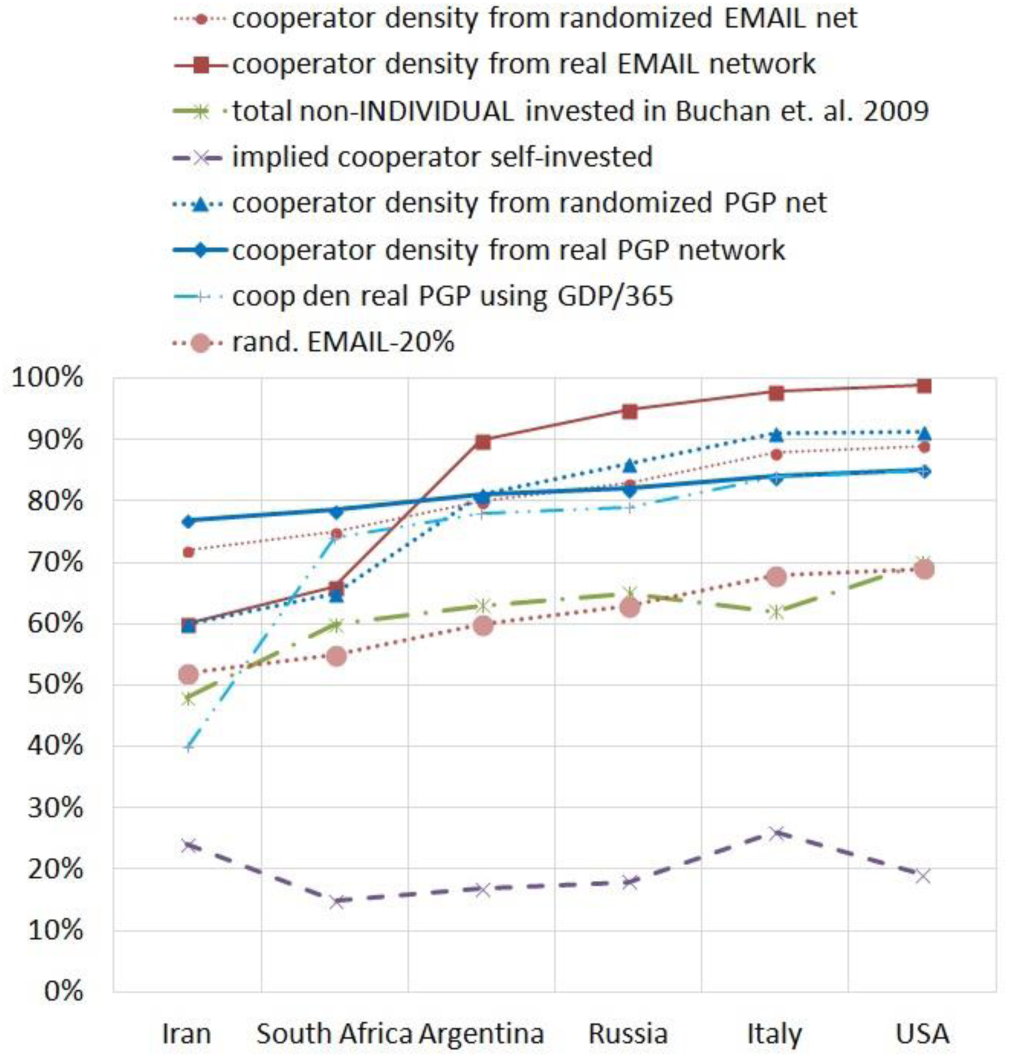
Percentage of other vs. self-investment from Buchan et. al. 2009 (investment) and Lozano et. al. 2008 (adjusted using GDP-relative payoff values compared to temptation)

The six countries are arranged as Buchan et. al. believed reflected their degree of globalization and interconnectedness, from low on the left to high on the right. The possible discrepancy that needs to be reconciled is that cooperation appears to increase as the size of connected networks increases to the right. From the simulated data on large and highly connected networks this is unexpected. If it cannot be explained, then there is a significant weakness in the simulation models, since we take the live-subject experiment to be the reference.

## Reconciliation Method

The problem may not only be the contextual incompleteness of the game definition. The players are not on a level playing field due to their wealth differences, which affect them personally and culturally.

Ito, Katsumata, Hasegawa and Yoshimura (2017) report that in chicken games (hawk-dove and snowdrift) it is *optimal* for the poor to cooperate more frequently, but not in Prisoner’s Dilemma (used by Lozano et. al.) or stag hunt, essentially because in Ito’s analysis a wealth (fitness) parameter *w* is *accumulated*. Accumulated geometric returns are damaged too greatly by relatively greater losses (compared to existing wealth) for the poor when making risky decisions. This character is peculiar to the chicken games. However, the principle that strategy may depend in part on a player parameter, not only on the arithmetical differences in game payoffs, means both that game payoffs are no longer independent of scale, and that strategy analysis made assuming the payoffs were independent of scale must be re-examined. Ito et al.’s method is a well-grounded departure from elementary game theory, though it has known counterparts in economic games. For example, in a price war the player with greater stored wealth wins.

Buchan et. al. use a multi-player game (twelve players in three groups) with multiple (three) and approximately continuous (ten tokens to be divided) investment choices, probably more accurately representing real cooperation problems with continuous investment choices. (Killingback and Doebeli 2002). The risk profile of this game is very different for self-investment than for other-investment (other being LOCAL or WORLD accounts, essentially a lottery depending on what eleven other participants, eight from different cultures, choose). Self-investment is a certainty while other-investment risks up to a 75% loss. This makes the Buchan et. al. game like the chicken games in a particular way. The poor should make the low-risk choice. Except in this case the low-risk choice is self-investment. If we think of other-investment as cooperation and the poor choosing cooperation, we would be misunderstanding the relatedness of the games. Their common point is risk rather than cooperation.

In Ezaki, Horita, Takezawa and Masuda (2016), who study reinforcement learning, it is found that [real] people consider whether their payoff (in cooperation or defection) is positive or negative based on whether a priori aspirations are met, rather than its absolute value. In realistic cases, we would assume aspirations would be a proportional change to one’s current wealth or fitness. This finding supports Ito et. al.

In the study of crash rate as a function of economic value-risk tradeoff (Shuler 2015 A) it has been found that to explain behavior such as a factor of three difference in per capita motorway death rates between otherwise similar countries (USA and Germany), it is necessary to consider the distribution of wealth within each country and separately calculate how each segment of the population responds to a perceived tradeoff (Shuler 2015 B). This is also supportive of wealth-relative decision making, even when fatality is one of the outcomes and the decision is personal.

It seems reasonable, based on these three considerations, that we should consider the payoff values for each investment strategy in Buchan et. al. relative to the wealth of the participants. We only have the country origin of the participants as a clue, but this may be enough as there are large differences in the countries. Figure 3 shows the per capita Gross Domestic Product (GDP) for each participant country for 2008 (the year preceding publication of Buchan et. al., in which we assume the experiment was performed, or thereabouts), which we will take as a normalization basis.

**Fig 3.**
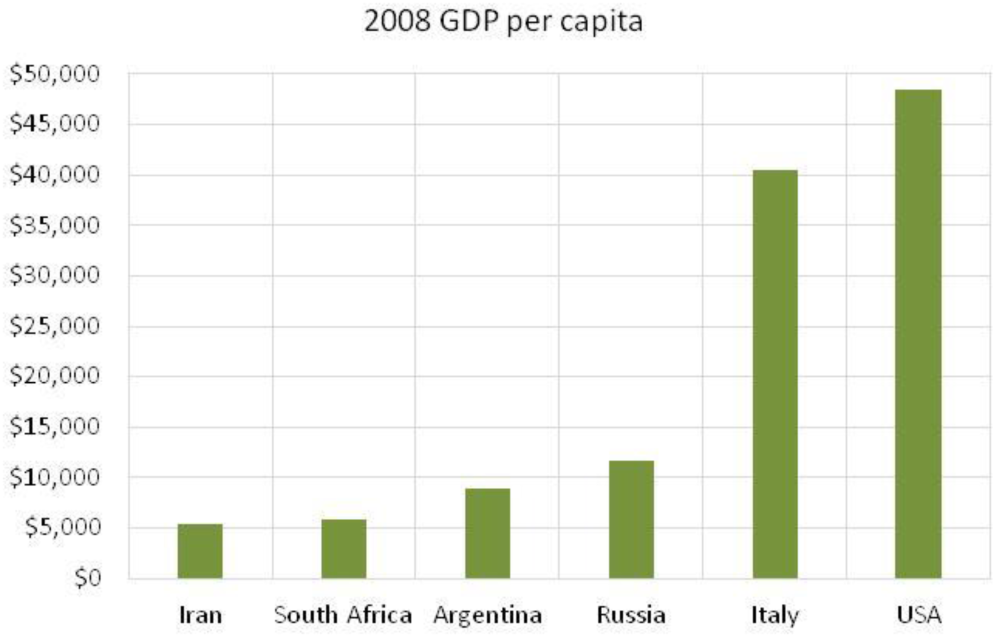
Prior year per capita GDP of countries in Buchan et. al. 2009.

There are two factors we don’t know in making such a scaling. First is the threshold of what value, relative to the country, a participant considers significant enough that it should not be placed at risk even for a possibility of larger reward of a given size. In the USA $15 is an urban lunch, whereas in Iran at the time it was a day’s expenses approximately. It is not the purpose of this paper to consider a methodology for accurately determining such scale factors, only to suggest that the scaling approach should be followed, so the author chose one that worked, basically. If the country GDP is divided by the number of days in a year, temptation in the left side countries seemed excessively high, close to *b=2* (see red dashed line in Figure 4).

**Fig. 4.**
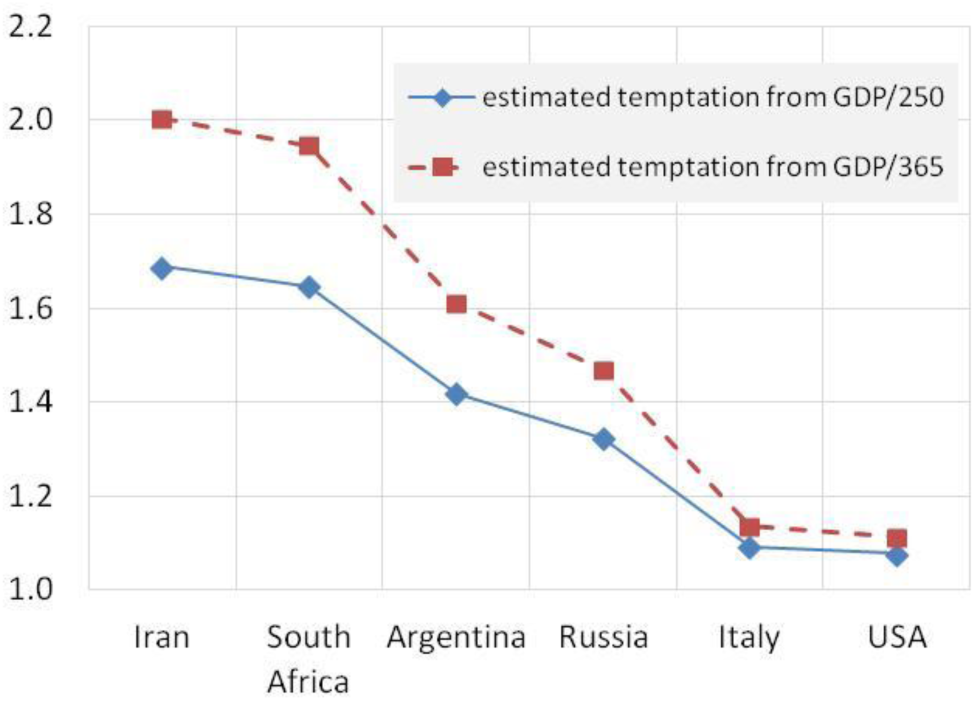
Estimated temptation as GDP/250 (workday) and GDP/125 (two workdays) normalization of $15 for each country in 2008.

If the country per capita GDP is divided by the number of work days, about 250, then the numbers correspond roughly to a day’s wages.

The other participants were strangers and most were not visible. It has been shown that humans use facial cues even in one-shot games in choosing strategies (Eisenbruch, Grillot, Maestripieri and Roney 2016), so they remained strangers.

Using the GDP/250 normalized temptation values, i.e. temptation to keep as a sure thing the $15, they are found to range between 1.06 and 1.55 as shown in Figure 4. Each temptation value is used with the Lozano et. al. data in Figure 1 to produce a density of cooperators estimate plotted in Figure 2. All four of Lozano’s temptation curves are used with GDP/250, but only one of them with GDP/365 for comparison. We will discuss these in a moment.

The second thing we don’t know is what network model to use. While the randomized network has less clustering, personal EMAIL networks may involve more discretionary choice of connections than economic networks. The more clustered real PGP network fails dramatically at high temptation. We might expect an economic network to fail less dramatically because it contains an element of necessity. One might also suppose that interaction with strangers and un-clustered networks trigger similar heuristic behavior, less cooperation in small things but not declining as fast at greater temptation.

Lastly, we ignore the difference in games and assume that participants may not even appreciate the particularities of the games, and may be mistaken at first (Burton-Chellew, Nax and West 2015).

## Results for reconciliation

Based on correlation of each candidate prediction from the Lozano simulation data weighted relative to country wealth (computed with Excel, see Table 1), the real PGP network weighted at GDP/365 has the best correlation, though all correlations are fairly good, ranging from 0.82 to 0.94.

**Table 1.** Correlation of cooperation density predictions from Lozano et. al. temptation data GDP/250 weighted except as noted

|  |  |
| --- | --- |
| non-INDIV | 1.00 |
| rand. EMAIL | 0.82 |
| real EMAIL | 0.84 |
| rand. PGP | 0.84 |
| real PGP | 0.85 |
| real PGP 365 | 0.94 |

Based on visual estimation the randomized EMAIL network from Lozano et. al. (2008) seems to be a reasonable predictive model for the Buchan study. In general visual estimation disagrees with the correlation data, with real PGP network weighted at GDP/365 appearing unsound for broad-spectrum prediction. There could be several reasons for this, principally the small number of country data points do not make a good statistical data set. The real PGP network has a rather violent discontinuity above temptation of 1.95, and the choice of GDP/365 positions one country just 0.05 beyond that. The direction of change between the first and second country is then correct but the magnitude far too great.

Moreover, the real PGP network is generally not a good choice not only because it may be more specialized, but because it has a sensitive dependence on calibration with respect to its near-discontinuity. Chances appear good a target experimental network will not have this discontinuity. The real EMAIL network did not, and the temptation-cooperation relations in Xu, et. al. did not, though those were artificial networks. The presence of discontinuities in the temptation-cooperation relations of “real” networks is an interesting question for further examination, as the presence of such discontinuities presents a problem both for forecasting and for the stability with respect to change of real networks.

Taking a linear fit approach, the overall slope of the randomized EMAIL network cooperator density function using GDP-per-capita/250 matches the slope of Buchan’s actual data from live subjects, with two countries (Iran and Italy) being slightly off the linear approximation, but not so far as to be unexpected and probably due either to other cultural factors, or sampling randomness. In the author’s judgment it is likely to have broader applicability, unless the target network is demonstrated to have discontinuities in its temptation-cooperation relation. The source of such discontinuities is likely to be extremely regular mid-scale structure, so that cooperation breaks down everywhere at once.

The difference between the Buchan et. al. data and our Lozano et. al. plus GDP derived data based on the randomized EMAIL network is shown in the dotted line at the bottom of Figure 2. This could be considered a second scale factor, a displacement constant. Whether it represents an error in predicting the density of cooperators (it might be the first time game theory models have over-predicted cooperation), or an amount withheld by the cooperators as suggested by the labeling in the figure, can be determined with additional data.

## Simulation of wealth-relative effects

It is important to understand how to produce wealth-relative effects in simulation, if as our analysis suggests they are present in real data. It is also important to understand how they appear in a game such as Prisoner’s Dilemma on which the temptation data was based. One clue we have already discovered is that wealth effects may be risk behavior rather than cooperation per se.

For these purposes a simulation was created using a tailored version of Prisoner’s Dilemma which we call the Farmer’s Game. It uses a payoff matrix similar to the one shown in Figure 5, and for the simulation network a lattice was used. The parameter “game cost-per-turn” effectively scales the entire payoff matrix for convenience (i.e.: add the game cost to each element).

**Fig. 5.**
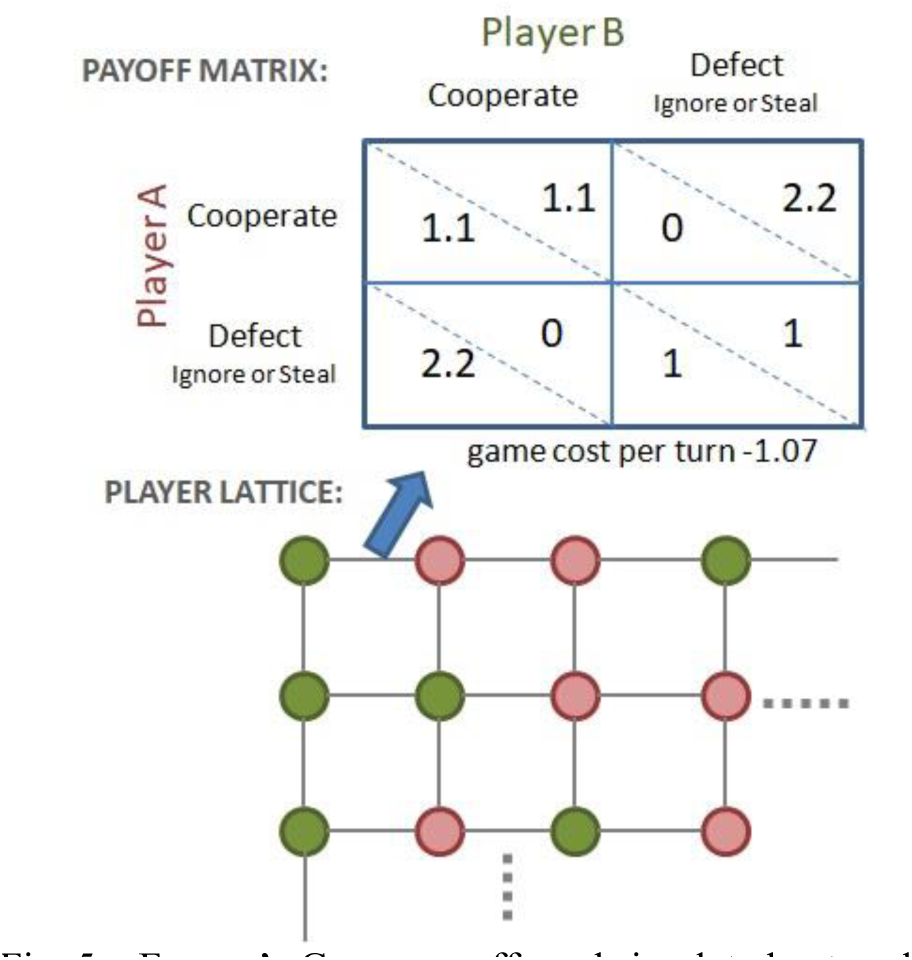
Farmer’s Game payoffs and simulated network.

The background for Farmer’s Game is that players will utilize the land between them to hunt, gather or grow food or produce other fitness-related items. If players ignore each other one token or “fitness unit” (or “food unit” if one prefers) is produced by each player as the mutual defection payoff.

If players agree to cooperate and “farm” the land, then an additional yield is available, which in Figure 5 is 10% with each player receiving a payoff of 1.1. Farming exposes both players to the risk that the other player may enter the field a few nights before the harvest and take more than his or her share. In the figure, the unilateral defection payoff of 2.2 indicates all the harvest is taken. Variations of these parameters will be considered. Note that while social dilemma games cannot be zero-sum, in most of our test cases any pair of payoffs in which at least one party is a cooperator in this game is conserved. Defection against a cooperator redistributes the benefit of cooperation, but does not remove it unless the defection is mutual. One conventional case will be demonstrated below in which the Defect-Cooperate move reduces total payoff. We might suppose the harvest is not mature or the defector loses some of it in haste.

Losing such a transaction is more than just a matter of scorekeeping due to three additional rules:

1. Players accumulate fitness (or food or wealth, etc.) proportional to their game payoffs as in Ito, et. al.
2. A player dropping to or below zero store of wealth or fitness tokens dies and is removed from the game and not replaced, so that we can track performance of initially assigned wealth and strategies.
3. The players must eat (or otherwise consume resources), as indicated by the game cost-per-turn, -1.07 in the figure.

It is easy to see most payoff matrices with simple integer payoffs quickly run away either negative or positive and do not remain near a death threshold, even if a game had one. So of course it only matters the relative performance between players, and a run of bad choices may be turned around (though most evolutionary research simply replaces a bad strategy and continues). Evolutionary approaches may not even produce all strategies of interest.

Thus in our simulation players do not update strategies (though some strategy elements will be dependent on wealth, which is updated), and dead players are not replaced. Adjacent players are instead connected to each other for further play. There is no wraparound and being on an edge can result in slower wealth growth or decay due to fewer turns per round. All of the following results use a lattice size of 20 × 20, which has a 20% edge population. A set of Monte Carlo simulations on a lattice of 100 × 100 produced no significant changes.

Wealth is initially randomly distributed with a parameter for density of the poor. Initial poor were given 4 tokens, approximately enough to last one turn with each neighbor, and wealthy were given 10. One simulation was conducted with 8 and 20, which essentially behaves as if everyone is wealthy.

In each round (or generation) a turn is taken with each neighbor. Most effects were evident after 10 rounds and had run their course by 20 rounds. All the results below were generated using 50 rounds x 4 turns each (except for edge players). Some results will show effects not only dependent on wealth, but on the magnitude of the cooperation excess return (10% in the figure for a

1.1 cooperation payout). This suggests another possible means of calibrating results to experimental situations based on cooperation payout. For example, in equity markets, long term mean annual return is in the neighborhood of 7.4% to 12.6% (Mehra and Prescott 2008). We did not explore this further at this time.

Either 3 or 5 simple strategies were used depending on the simulation. These were

1**.TitTat** – Classic tit for tat with 10% forgiveness (variable by parameter). This was implemented as a memory one strategy with each neighbor. In addition, TitTat serves as a default strategy for all the others if their special conditions are not met.

2**.Subsist** – If survival of the next round (of four turns) is seriously in doubt (present wealth 4 or less), then defect, else use TitTat strategy.

3**.Exploit** – If at least 2 times as wealthy as other player, defect, else TitTat.

4**.Thief** – If the other player is at least 2 times as wealthy then defect, else TitTat.

5**.Middle** – If survival of 2 rounds in doubt (present wealth 8 or less) then defect, else TitTat.

The primary objective is to see if and under what conditions the Subsist strategy has an advantage, as well as to generally record the history (population and wealth) of each category of players. The Exploit strategy is primarily included to create some danger in the simulation, otherwise only the poorest players would ever defect. The Middle strategy is a more conservative version of Subsist, intended in some way to represent the middle classes, though this is not verified. While the Middle and Subsist strategies are more risk averse than TitTat, the Thief strategy is more risk aggressive, attacking wealthier players.

An overview of simulation results with respect to variation of density of the poor and magnitude of initial wealth, otherwise at the default parameters of Figure 5, is given in Figure 6 below. We will discuss those default parameters in connection with equilibrium.

**Fig. 6.**
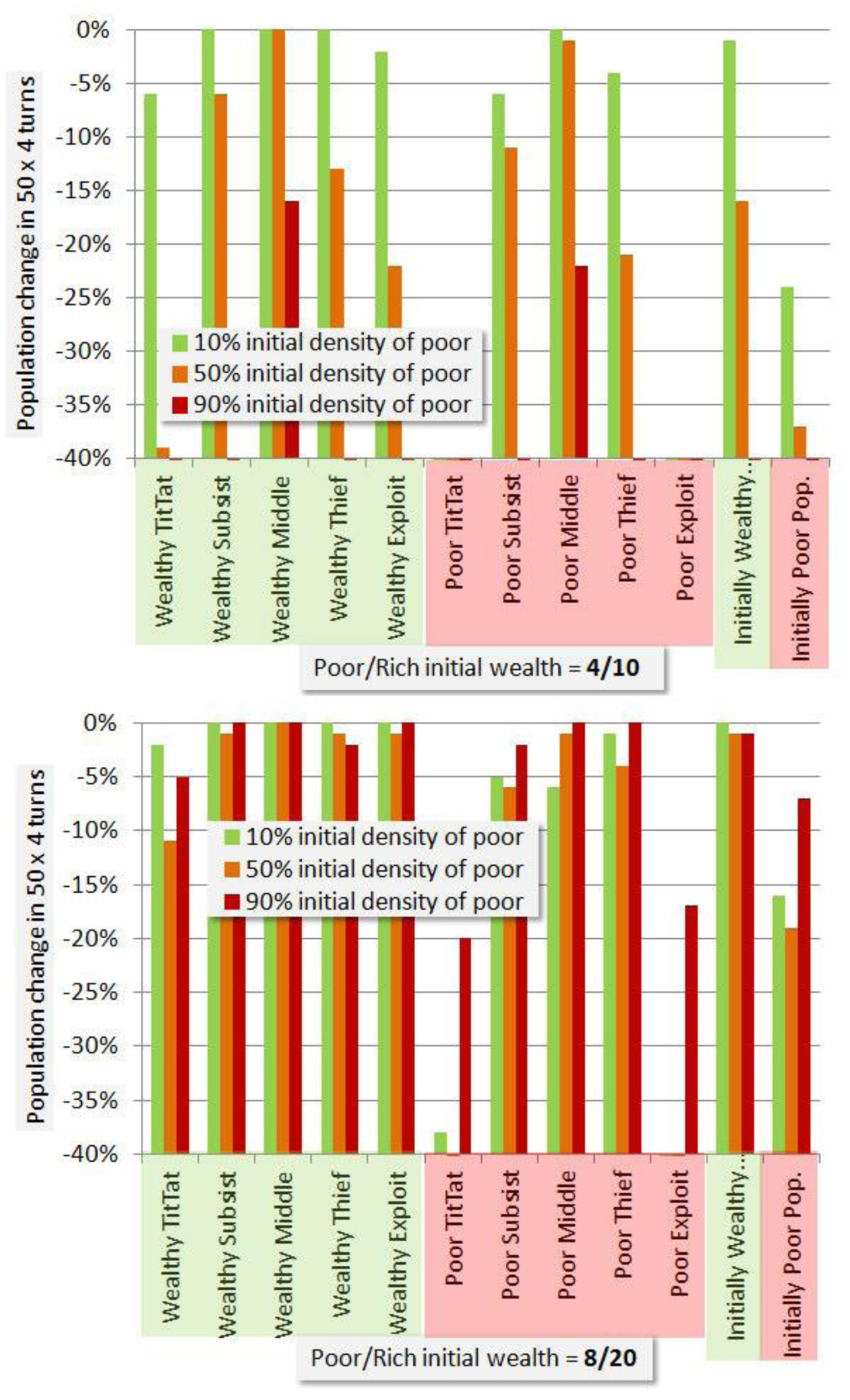
Simulated effect of variation in poor density and initial wealth with cost-per-turn of 1.07.

There are several interesting points in this result. The initial poor/rich values of 4/10 are in a good range to show wealth effects as expected. The poor do somewhat worse than the wealthy, and the wealthy do worse if the initial density of poor is high. TitTat, is evidently the worst strategy for the wealthy in this mix. At 90% initial poor distribution, only users of the most conservative strategy, Middle, are surviving. In the 8/20 poor/rich mix, all the wealthy categories do fairly well, though again TitTat not as well as the others. For the poor, TitTat and Exploit are wiped out except for the 90% case in which the poor are mostly playing against other poor.

Keep in mind when viewing these results that they are classified with respect to initial wealth. While some players die off, or become poor, a number of players become wealthy in most simulations. However, identifying results based on final status obscures the history we are looking for. All results are based on 100 simulations at the given parameters with rich/poor status and strategy distributed randomly in each, and results averaged, using a 20 × 20 network lattice. Below we will give histograms.

Figure 7 shows sensitivity to TitTat forgiveness. This affects all strategies, since all use TitTat is their base. The 10% forgiveness rate we have selected as our baseline gives better performance than 5% or 50%. While it is better for Thief and Exploit and Subsist as might be expected, more surprising is that it is better for everyone including TitTat, being the only condition under which any pure TitTat strategies of any wealth survive 50 rounds (∼200 turns).

**Fig. 7.**
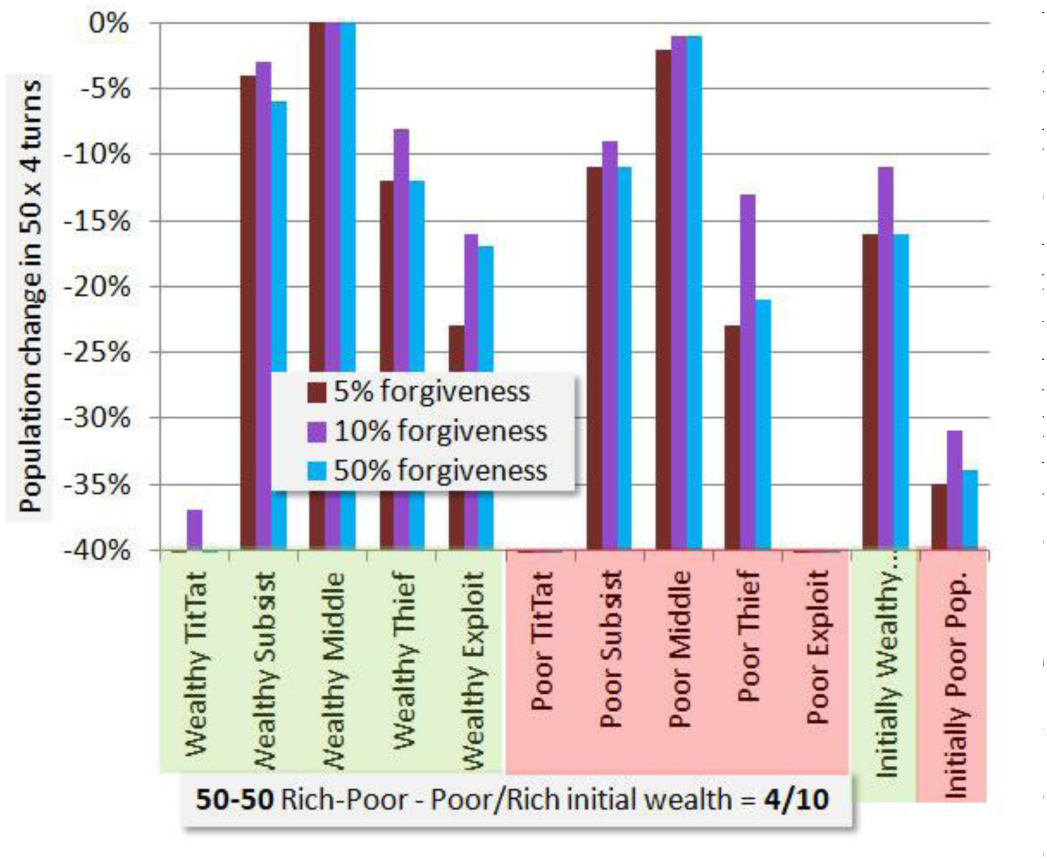
Simulation sensitivity to TitTat forgiveness.

It appears that optimizing forgiveness, for this strategy mix, is not a social dilemma. A value which approximately optimizes the fate of a decision maker also benefits all others in terms of survival rate. Presumably wealth is less concentrated, as the forgiveness effectively shares wealth with Subsist, Middle, Thief and Exploit on occasion, so if survival is not the dominant motive there may still be a social dilemma. There is no moral justification for Exploit, since it is theft by the rich. Subsist and Thief are means for the poor to get by, to escape the apparent extinction of poor TitTat players.

In Figure 8 we explore softening the impact of the Defect-Cooperate pair of moves. In addition the fourth payoff combination is the promised conventional Prisoner’s Dilemma payoff in which some social benefit is lost by the Defect-Cooperate pair. In this case (2.0|0|-1.05) the equilibrium for this payoff of -1.05 cost-per-turn is used. It shows a performance similar to (2.2|0|-1.07), but just slightly worse.

**Fig. 8.**
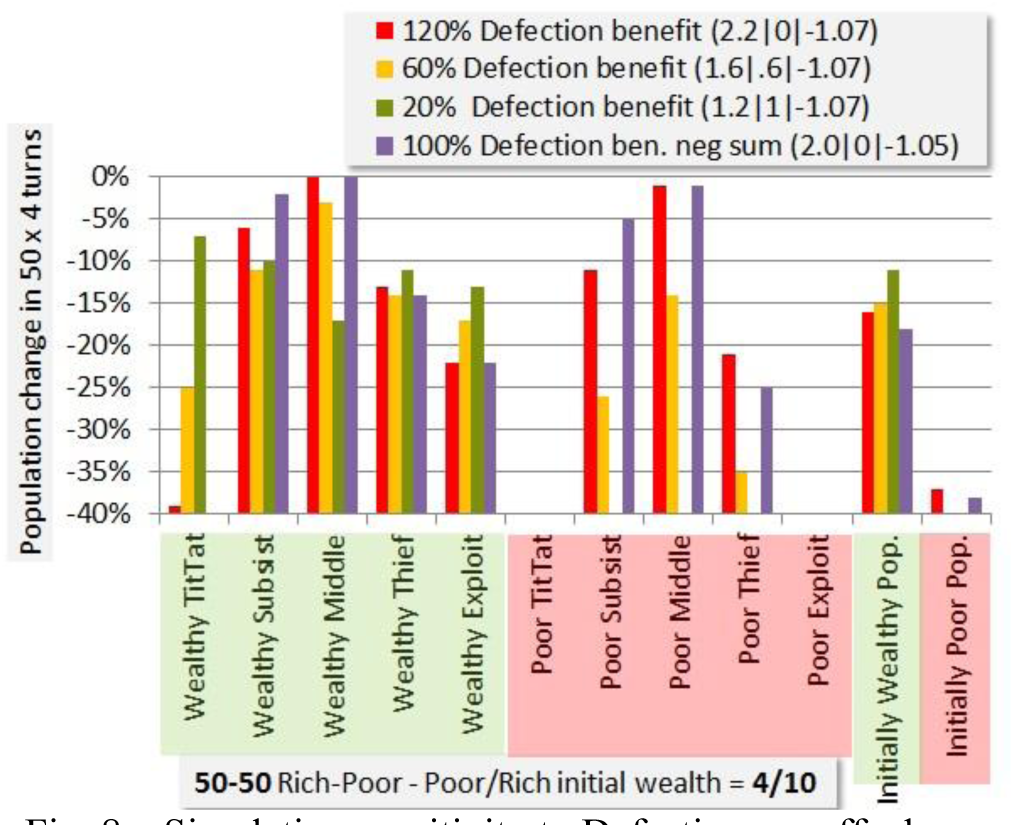
Simulation sensitivity to Defection payoff when other player Cooperates.

In principle there is no motive for the Defector to take less than all of the harvest, as a TitTat player (which all of them are in base mode) will note one defection and present the same magnitude of response in any case. However in real life other more severe penalties may deter full defection, so we compare the baseline 2.2|0 payoff combination for D|C to 1.6|0.6 and 1.2|1. Players in this simulation were not “aware” of the payoff magnitude being changed, so this does not measure temptation to defect, which is fixed in the heuristic strategies, but changes the wealth distribution and therefore survival consequences.

There are two very striking consequences. The TitTat strategy is “rescued” by the lower impact of a partner defecting. Unfortunately this devastates the poor who depend upon the defections for survival.

These findings already justify our hypothesis that with a death threshold and realistic cooperation benefit (not growing or declining in wealth rapidly away from the threshold) that wealth effects are evident in a game payoff matrix that qualifies as a version of Prisoner’s Dilemma. But the move choice that is favored is the one which is low risk, defection, not necessarily cooperation.

We now find the wealth equilibrium cost-per-turn using the 4/10 wealth split and 50-50 wealthy-poor distribution. The three strategies TitTat, Subsist and Exploit are evaluated separately from the 5-strategy mix, with results shown in Figure 9.

**Fig. 9.**
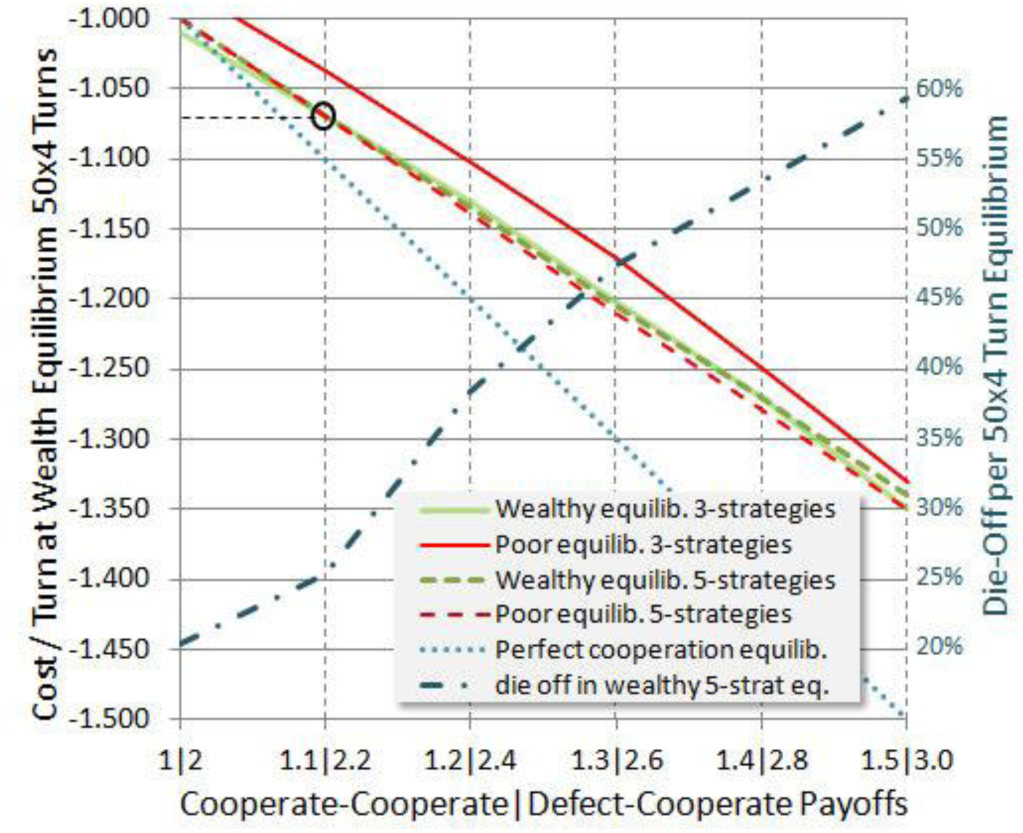
Wealth equilibrium cost-per-turn.

This analysis finds total wealth equilibrium separately for initially rich and initially poor players, for the two strategy groupings. Perfect cooperation is represented by the dotted blue line, as if all players were TitTat with no noise. In this case no player dies, however the equilibrium turn cost matches the entire benefit from cooperation.

Societies that we experience tend to exist somewhere close to equilibrium, changing slowly over time unless there is some external trigger. This means that by finding equilibrium we have a clue to an operating point. The reasons are many. Costs or conflicts might rise as populations increase or resources become scarce. Even development of new technology is a cost, one with which many are now engaged in demanding jobs. One may also view the turn cost as the standard of living, in which case a higher turn cost is not necessarily negative if the members of society can afford it.

It appears that 1.07 is the equilibrium cost-per-turn for three of the four categories, with only the poor in the less diversified 3-strategy mix having a slightly lower equilibrium. In that sense, the data of Figure 6 put the 4/10 poor at a disadvantage, since Figure 6 was based on the 1.07 equilibrium. However one can interpret that as just another wealth-relative effect: that the poor cannot keep up in a society that operates near an equilibrium standard of living, expecting certain housing and transportation and healthcare and occupational safety and so forth standards of everyone.

On the right side of Figure 9 is a scale for reading the die-off rate of one category. Die-off rates are similar for categories that have a similar equilibrium value. Perhaps the most disturbing thing about higher cooperation payoffs (benefits) is that they entail highly elevated die-off levels (or bankruptcy levels, if one is considering this to be only a financial game), unless one assumes perfect cooperation. In the author’s view perfect cooperation is desirable but unlikely, and we should grapple with the reality. Evolutionary studies are needed to ascertain the likelihood, or not, of every obtaining perfect cooperation, which we do not address here. We only evaluate mixes of given strategies, and specifically avoid highly tuned or optimized versions of them for generality.

Network architectures and degree of persistence in interaction are known to have large effects on strategy evolution. Without evolution we do not expect great deviation for other networks. Though persistence is high in our model, the Monte Carlo runs average a tremendous variety of configurations, such as groupings of cooperators, or lone cooperators surrounded by Subsist or Thief players. A histogram of the 3-strategy survival rates is shown in Figure 10.

**Fig. 10.**
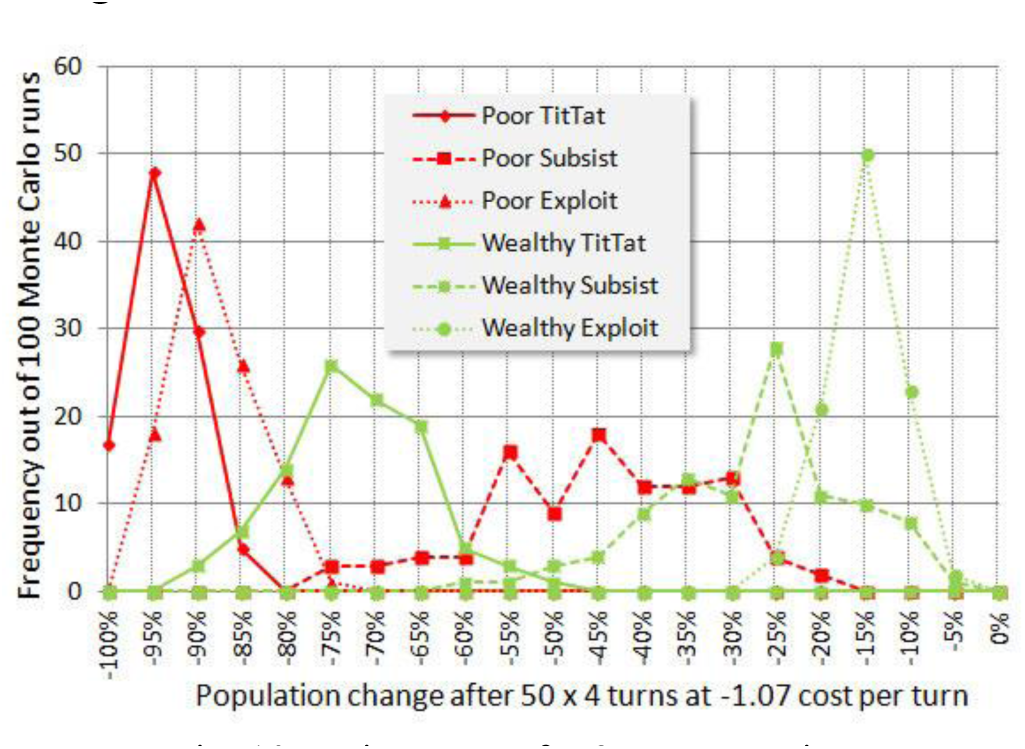
Histograms for 3-strategy mix.

Four of the histograms are quite close to normal distributions. The two Subsist categories have a high variance and are only loosely normal. They become normal at 300 or more Monte Carlo iterations, but their average value appears usable at 100 iterations, facilitating the examination of a large number of cases.

Figure 11 shows a different way of visualizing the variation of simulation performance, showing survival rates for each category vs. cost-per-turn for the baseline 1.1|2.2 mutual-cooperation | defection-if-cooperating payoff. The 3-strategy mix has evidently a different character, performing somewhat less well than the 5-strategy mix, but having a more forgiving slope as cost-per-turn increases.

**Fig. 11.**
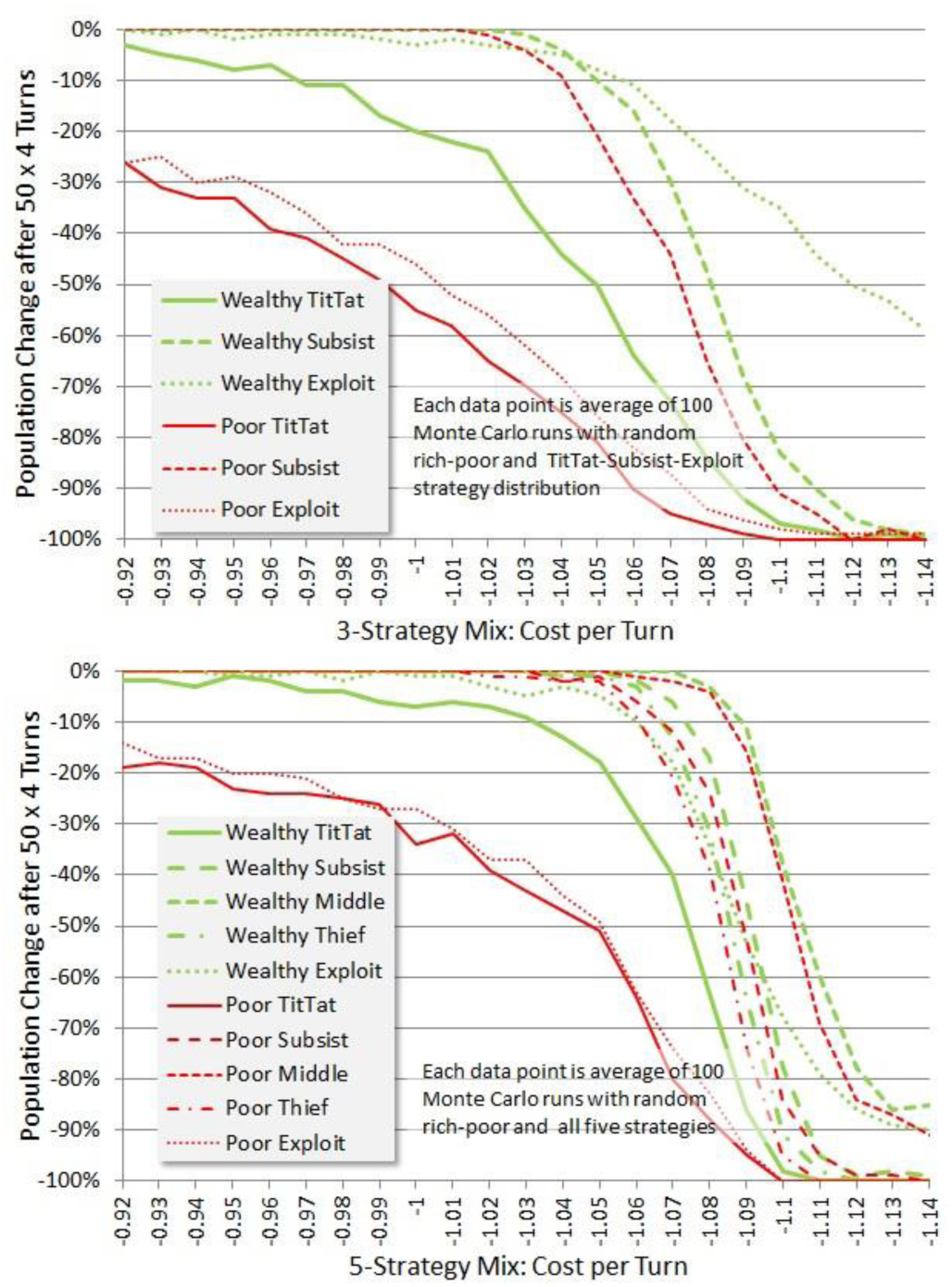
Survival vs. Cost-per-Turn at 1.1 cooperate payoff.

The equilibrium of the poor in the 5-strategy mix has come up because the Middle and Subsist strategies perform so well, though other poor player strategies do somewhat better. We might tentatively ascribe a benefit to diversity of strategies, perhaps due to less likelihood of unfavorable clusters forming (which might or might not hold up in an evolutionary simulation). But as the cost/standard of living increases it may be more prone to abrupt failure. And it may be very important to explore this further as a hypothesis in an experimental setting.

The wealth-relative effect remains clear in Figure 11, and seems to be attributable specifically to two sources. TitTat is a poor strategy for the poor if it entails risk. They must take no chances with survival. And as badly as wealthy TitTat does, poor TitTat does much worse. Second, the poor cannot afford to use risky or rarely applicable strategies such as Exploit. In an evolutionary simulation, we might expect these players to be eliminated and replaced by more savvy players (whether intentionally or heuristically savvy). One wouldn’t necessarily expect a wealth effect to disappear in an evolutionary simulation, as wealthy players often become poor in simulations, and may find their strategies maladapted.

Stewart and Plotkin (2014) report finding that in co-evolution of strategies and payoffs (via iterated Prisoner’s Dilemma) that the benefits of cooperation may be pushed higher, and that tradeoff with the cost of cooperation may lead to a dramatic collapse. A trend toward increases in benefits in cooperation leads via the evidence of Figure 9 to a higher die-off rate at equilibrium, toward which our own arguments have assumed society will be pushed. A high die-off rate (60% over the 50 rounds at a benefit of 1.5, though we do not know exactly what time period this might correspond to) of known relationships is at least not inconsistent with Stewart and Plotkin. At the 1.5 cooperation benefit equilibrium is 1.35 and the population change shown in Figure 12 though not as sharp as at 1.1 benefit is in heavy die-off.

**Fig. 12.**
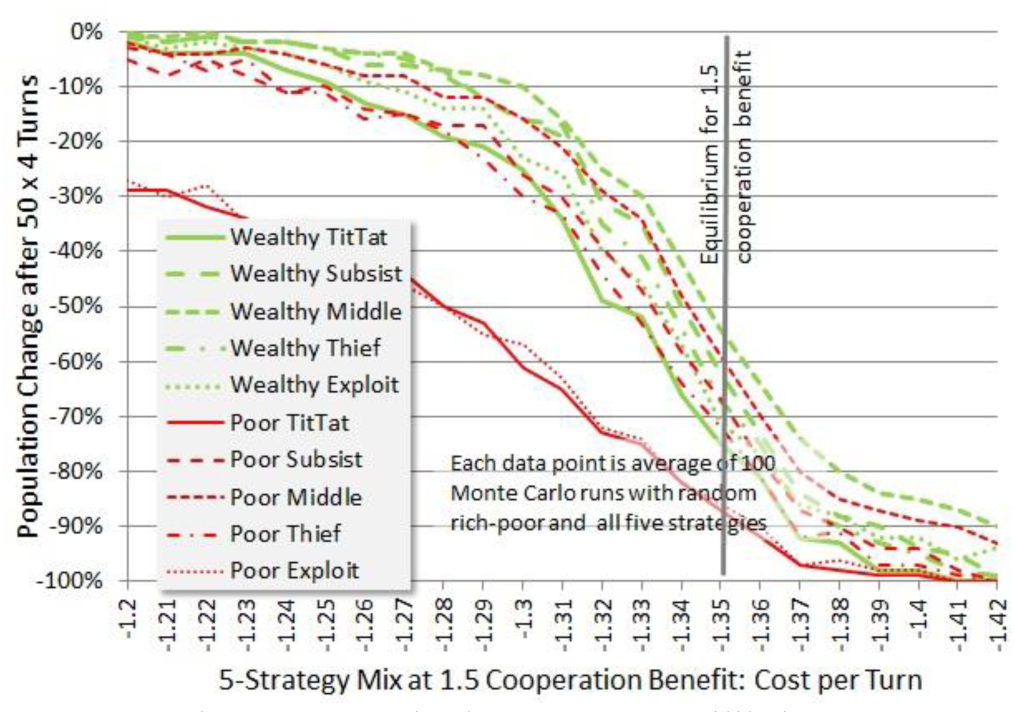
Survival near 1.35 equilibrium for 1.5 cooperation benefit.

Our final view of the data in Figure 13 shows wealth (rather than population) by strategy and initial wealth category for the baseline 1.1 cooperation payoff as cost-per-turn varies.

**Fig 13.**
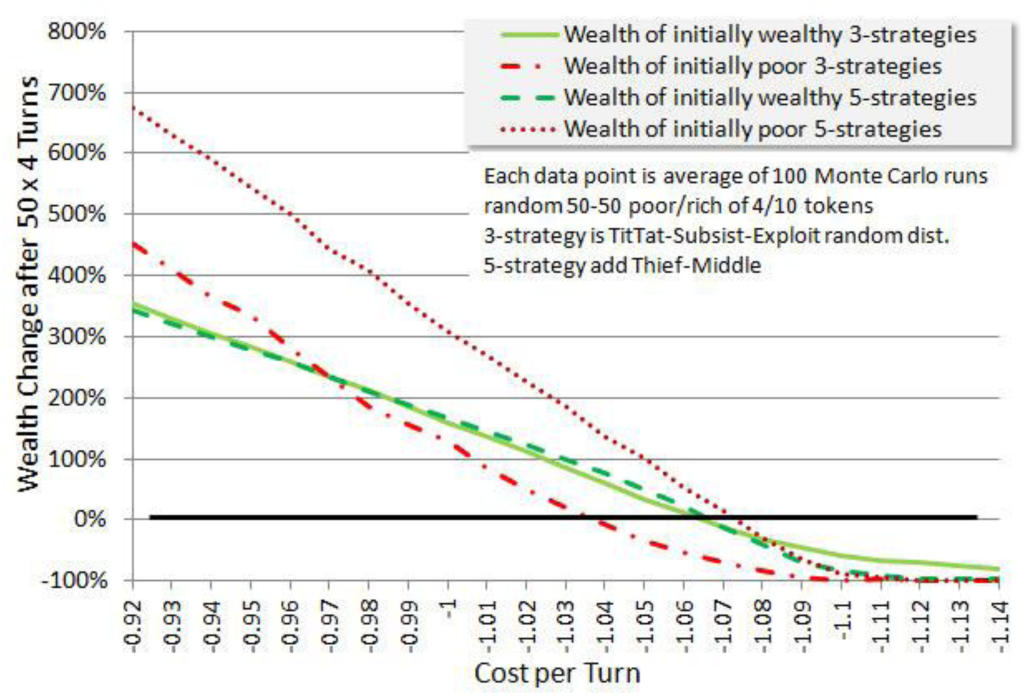
Wealth after 50 rounds by category.

Figure 13 reveals the “up side” of being poor, in our simulated world, is a small base for measuring future wealth increases, and for any case of cost-per-turn below their equilibrium value the poor get richer faster than the wealthy as a result of this comparative artifact. However, before too much optimism is imbued, note that in the “real world” wealthy “farmers” are not constrained to invest the same 1.07 tokens per turn as everyone else in the game, but may expand their reach in geography and industry to grow wealth exponentially, as ordinarily assumed in finance.

A second reason for presenting Figure 12 is to graphically illustrate just how fast wealth can increase even in the linear fixed-payoff game case, making the poor rich, and moving everyone in a simulation far from the survival threshold. Except for carefully tuned payoffs, everyone either becomes wealthy or dies. This is more than just a reason for lack of note of wealth-relative effects in most simulations. It is importantly a reason why we argue most societies we observe are in fact operating close to equilibrium. Those which do not either die off, or become wealthy and out of the reach of study by social scientists, flying around in private jets avoiding airport security and cooperation studies.

## Discussion

The experiment conducted by Buchan et. al. is extremely important. The results of our paper do not contradict their findings, which are empirical, or their reasoning that these findings are likely due to globalization. However, there is more than one way in which they could depend on globalization. They could be due to the different connectivity in the globalized countries, or due to the greater affluence of those countries which may well be in part due to globalization, in which case all citizens in the participating countries need to share in the benefits or the level of cooperation may not be maintained.

Other areas of social change exhibit the same kind of ambiguity between social network and affluence as cause or effect. Sander and Putnam (2010) point out that in the early 1960s more than half of all Americans said they trusted others, but it was only one third in 2010. Since 2001 only the upper-middle-class young people have remained civically engaged while pay-for-play extracurricular activities and teaching-to-the-test have encouraged others to drop out. Can this be corrected by encouraging adults and youth to be more socially connected? Or do we have to address the economic mechanisms by which the affluence gap has widened as Putnam, Frederick and Snellman (2012) seem to suggest two years later?

The amount participants were willing to risk in the Buchan et. al. study is only a fraction of a day’s pay ranging from a tenth to a half, small when compared to the investment necessary to address real public goods issues such as climate change. Risk was an important element in our framing of the game to decide how to apply relative payoff value. This will vary with both game structure, and knowledge the participants have about each other, in this case very little. Apparently it also varies with network structure, since the randomized (de-clustered) network better fit the Buchan et. al. data. Are low clustering and stranger interaction both proxies for some kind of risk? Depending on simulation model parameters there likely are different effects, for example clusters of cooperators being “food” for defectors in basic scenarios. In memory models where tit-for-tat has evolved, there may be greater risk of retaliation from strangers. Either of these could be dependent on the magnitude of the temptation/defection, and explain the crossover in Figure 1 at high temptation where the randomized network becomes the higher cooperating one.

With Lozano et. al. partly confirmed by reconciliation with Buchan et. al.’s experimental data, the principle of relativity of temptation/payoff values introduced, and a possible relation between strangeness and network structure at the hypothesis stage, many interesting possibilities for investigation are apparent. We can postulate a model of a class-structured society in which individual cooperation depends upon the magnitude of the payoffs of each “game.” Would such a model show classes cooperating with their peers, but taking risk-averse strategies (whether they be cooperation or defection) when cooperating with more wealthy classes and vice versa? If in such a model one friend or relative becomes wealthier than another, would the dry mathematical model produce a change in cooperative behavior we might call “envy”? Do class structures play a role in evolutionary stability?

## Conclusion

It appears that we can conclude that minimizing the risk of catastrophic loss, though only when it manifests in some near timeframe (to be determined by context and future research, suggested by the success of the Subsist and Middle strategies), is advantageous, at least among simple competing strategies as in the simulated examples. Other interesting wealth-related effects are present such as the unexpected effect of a high density of poor on the rich. The effects are noticed not only for chicken games, but to variations of Prisoners Dilemma with appropriate wealth properties, which we called a Farmer’s Game. A single paper can hardly scratch the surface of all the possibilities, and further investigation is needed.

Relative payoff value, when risk differences are evident between strategies, would appear to be useful for understanding the relation between highly conceptual (theoretical) or simulated results and the behavior of actual humans either in experiments or in-situ. This sort of practical experimental confirmation increases confidence when applying cooperation theory to the development of specific policies and societal goals. Some players can endure a long string of losses and others can’t. Asking them to do so may induce affluence changes which increase class differences, disrupt social networks and degrade cooperation; changes which are hard to reverse because of the ambiguous mutual causality of affluence and cooperative social networks. Even if the losing players still seem well off to outsiders, their aspiration-relative view of the decreasing payoffs is negative.

Addressing wealth-relative effects in an evolutionary simulation, especially as not only strategies and population numbers will evolve, but also the payoff and cost parameters, may be quite complex. The hope embodied in this paper is that sufficient introduction of the topic has been made that some researchers will consider it, ever mindful of the utility of attempting to reconcile theoretical and simulation results with empirical experiment, which is also facilitated by the wealth-relative formulation.

Cooperation theory has made rapid advances since the early 1980s, and may now be able to be applied in the real world. If appropriately scaled and calibrated it may help safely and prudently “engineer our society” as Ezaki et. al. (2010) suggest. But as cooperation is very valuable, it is also subject to the misadventures of hasty pursuit.

## ACKNOWLEDGEMENTS

The author thanks B. G. Smith and T. Koukouvitis for numerous discussions and suggestions, and T. Koukouvitis for review of the final manuscript.

## SIMULATION AND DATA AVAILABILITY

The reader may find an interactive version of the simulation at http://mc1soft.com/papers/wealthrelative/ and/or archived at a suitable repository recommended by the journal of publication, along with all data from this paper in spreadsheet format.

